# Structural transition of the ground-state structure to steady-state structures by sequential binding of ATP to V/A-ATPase

**DOI:** 10.1101/2022.10.04.510906

**Authors:** Atsuko Nakanishi, Jun-ichi Kishikawa, Kaoru Mitsuoka, Ken Yokoyama

## Abstract

V/A-ATPase is a rotary ATPase that shares a common rotary catalytic mechanism with F_o_F_1_ ATP synthase. Structural images of V/A-ATPase obtained by single-particle cryo-electron microscopy (cryo-EM) during ATP hydrolysis identified several intermediates, revealing the rotary mechanism under steady-state conditions. Here, we identified the cryo-EM structures of V/A-ATPase corresponding to short-lived initial intermediates during the activation of the ground state structure by time-resolving snapshot analysis. These intermediate structures provide insights into how the ground-state structure changes to the active, steady state through the sequential binding of ATP to its three catalytic sites. All the intermediate structures of V/A-ATPase adopt the same asymmetric structure, whereas the three catalytic dimers adopt different conformations. This is significantly different from the initial activation process of F_o_F_1_, where the overall structure of the F_1_ domain changes during the transition from a pseudo-symmetric to a canonical asymmetric structure. Our findings will enhance the future prospects for the initial activation processes of the enzymes with dynamical information, which contains unknown intermediate structures in their functional pathway.

## Introduction

Vacuolar/archaeal-type ATPase (V/A-ATPase) is an excellent molecular machine that shares a common rotary catalytic mechanism with F_o_F_1_ ATP synthase [1-5]. V/A-ATPases are found in archaea, with some eubacteria species utilizing a transmembrane electrochemical ion gradient to synthesize/hydrolyze ATP [6-9]. V/A-ATPases are structurally and evolutionarily related to vacuolar-type ATPase (V-ATPase) [2, 9] but have evolved different regulatory mechanisms based on their function [5, 10-13].

The thermophilic bacterium *Thermus thermophilus* (*Tth*) possesses V/A-ATPase, which functions as an ATP synthase in *vivo* [6-8, 14]. V/A-ATPase has a simpler overall structure than the eukaryotic V-ATPase that has a subunit stoichiometry of A_3_B_3_D_1_E_2_F_1_G_2_*d*_1_*a*_1_*c*_12_ (Figure 1A, Figure 1-figure supplement 1) and has structural stability that allows for direct observation of the rotary motion in the enzyme by single-molecule experiments and X-ray crystallography of the subunits or subcomplexes [15-22].

**Figure 1.**
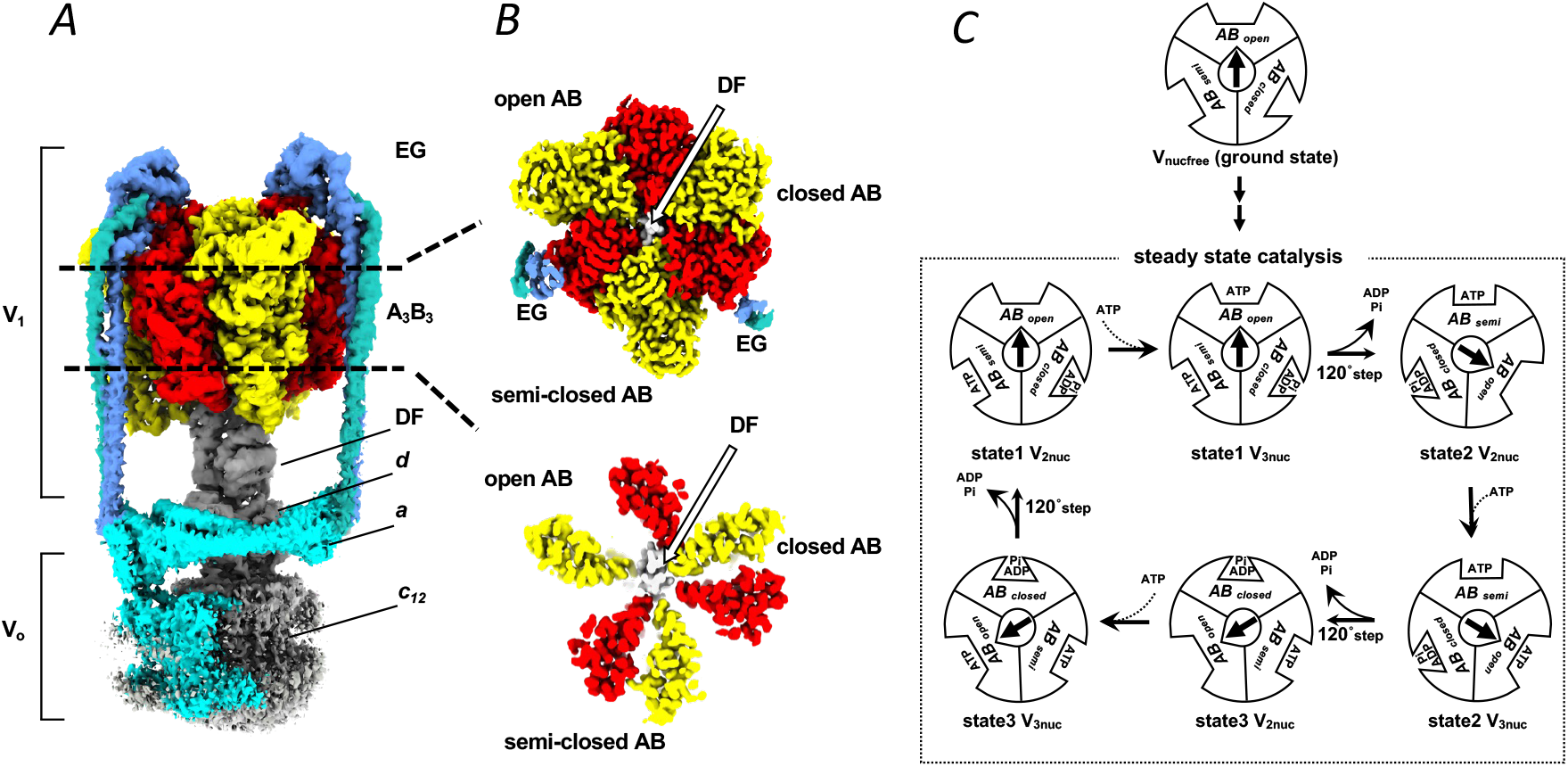
Cryo-EM maps of the V/A-ATPase and schematic representation of steady-state catalytic cycle driven by ATP hydrolysis. Side view of the representative map of the whole complex (**A**). Cross sections of the nucleotide-binding sites (top) and A_3_B_3_ C-terminal region (bottom) of V_2SO4_ map viewed from the top (**B**). **C**, Three rotational states (state1–3) interconverted through a tri-site mechanism where the site occupancy alternates between two and three as shown in V_2nuc_ and V_3nuc_, respectively. Schematic showing the three different catalytic states of AB subunits denoted AB_open_, AB_closed,_ and AB_semi_. As the rotor subunit (black arrow with bolt type) rotates 120°, three AB dimers altered to the next state; AB_closed_ to AB_open_, AB_open_ to AB_semi_, AB_closed_ to AB_open_ (e.g. state1 V_3nuc_→ state2 V_2nuc_). The activation process of nucleotide-depleted enzyme (ground state enzyme, upper) has not been elucidated.

In the V/A-ATPase, the V_1_ domain (A_3_B_3_D_1_F_1_) can function as an ATP-driven rotary motor, and sequential ATP hydrolysis by the three catalytic sites on the AB dimers produces conformational changes in the A_3_B_3_ hexamer, which induces a 120° rotation of the central DF rotor [6, 8, 13]. The catalytic cycle is often summarized as a circular reaction scheme that indicates catalytic events linked to the rotary position of the central axis (Figure 1C). Single-particle cryo-electron microscopy (cryo-EM) analysis of the V/A-ATPase identified three rotational states (state 1–3) and an asymmetric structure of the three AB dimers, termed “open (AB_open_),” “semi-closed (AB_semi_),” and “closed (AB_closed_) (Figure 1B). Even in the presence or absence of nucleotide binding, conformations of both AB_semi_ and AB_closed_ remained closed, which does not allow easy access of ATP to the catalytic sites [23, 24]. Therefore, ATP binding occurs in AB_open_ where the catalytic site is fully open. In contrast, a recent structural study of bacterial F_o_F_1_ using cryo-EM revealed that nucleotide-depleted F_o_F_1_ comprises three catalytic β subunits, all of which adopt an open form [25]. Thus, the accessibility of nucleotides to the three catalytic sites was significantly different between the F- and V/A-type ATPases in the absence of nucleotides.

Structural snapshots of V/A-ATPase under steady-state conditions were obtained using cryo-EM [23]. These structures revealed that all three catalytic sites on the AB dimers were occupied by ATP/ADP under the waiting conditions for catalysis ([ATP] > *K*_m_) in the ATPase cycle (Figure 1C, V_3nuc_ structure). Contrary, only the catalytic site on AB_open_ was empty (Figure 1C, V_2nuc_ structure) under the waiting conditions for ATP binding ([ATP] < *K*_m_). These results support the tri-site mechanism of V/A-ATPase, where site occupancy alternates between two and three [26-28]. Moreover, the structural snapshots indicate three catalytic processes simultaneously proceed; i) zipper movement in AB_open_ with the ATP, ii) ATP hydrolysis on AB_semi_, and iii) unzipper movement of AB_closed_ with the release of ADP and P*i*. In other words, simultaneous interlocking of the three catalytic processes and 120° rotation of the axis are necessary for continuous catalytic reactions [23].

Although steady-state intermediate structures of VA-ATPase were unveiled in a previous study using cryo-EM, how the three catalytic sites of the enzyme are filled with nucleotides in the initial activation process remains unresolved. Because the A_3_B_3_ hexamer of the nucleotide-free V/A-ATPase (V_nucfree_) also adopts an asymmetric conformation (AB_open_, AB_semi_, and AB_closed_), the first ATP likely binds to the catalytic site on AB_open_ in V_nucfree_. For ATP to bind to the following binding site, the conformation of AB_closed_ should be changed to AB_open_ with a 120° step of the rotor. However, it is unlikely that the 120° step of the rotor would occur only by the zippering motion of AB_open_ with ATP because simultaneous catalytic events at three catalytic sites are also required for the steady-state turnover of the V/A-ATPase. The measurement of ATPase activity of *Tth* V_1_-ATPases presents an initial lag with low ATPase activity and subsequent activation of the ATP hydrolysis reaction [29]. In addition, the lag time was significantly shortened as the ATP concentration increased in the reaction solution. However, the conformational change process of V/A-ATPase from the initial state, referred to as the ground state, to the steady state upon ATP binding remains unknown (Figure 1C).

In general, it is challenging to identify the initial catalytic state of the enzymes by canonical cryo-EM observation because the activation process is completed within a few seconds [30-32]. Thus, structures focusing on the initial reactions of rotary ATPases have not yet been obtained.

Here, we report cryo-EM observations to capture the short-lived intermediates of the V_1_ domain under different reaction conditions and time points following initiation by ATP with sulfate, which extends the duration of the initial inactivated state of V/A-ATPase. The captured sequential intermediate structures provide the first insight into how V/A-ATPase changes its conformation from the initial state to the steady state through the sequential binding of ATP to its three catalytic sites.

## Results

### Extension of the initial activation process of V/A-ATPase upon sulfate addition

In this study, we used the V/A-ATPase, including TSSA mutation (A/S232A, T235S), which is much less susceptible to ADP inhibition than the wild-type enzyme [8, 14]. The bound ADP was easily removed from V/A-ATPase by dialysis against phosphate buffer. We termed the V/A-ATPase without nucleotide V_nucfree_ (nucleotide-free V/A-ATPase) as the ground state structure. The resulting V_nucfree_ was reconstituted into a nanodisc and used for subsequent analysis.

The initial lag phase of the ATP hydrolysis profile by V_nucfree_ was observed, suggesting the existence of initial inactivated intermediates (Figure 2A, B). This lag phase disappeared within a few seconds under saturating ATP conditions (Figure 2A, control). The longer the activation time, the easier it is to capture short-lived intermediates using cryo-EM during the activation process for the initial inactivated V/A-ATPase. Therefore, sulfate ions, which are known to decrease the reaction rate as phosphate analogs, were added to the reaction mixtures to extend the initial lag phase [29]. Expectedly, the initial lag time was significantly extended in the presence of sulfate under saturated ATP conditions, which is suitable for capturing snapshots of intermediates for a specified time interval (Figure 2B). When comparing the sulfate concentration dependence on ATPase activity, it appears that the presence of 20 mM sulfate does not significantly affect the ATPase activity of the enzyme and only extends the duration of the initial lag (Figure 2A, Figure 2-figure supplement 1). In contrast, V/A-ATPase exhibited little ATPase activity at much lower ATP concentrations than the *K*_m_ value (a final concentration of 50 μM) in the presence of 20 mM sulfate (Figure 2C).

**Figure 2.**
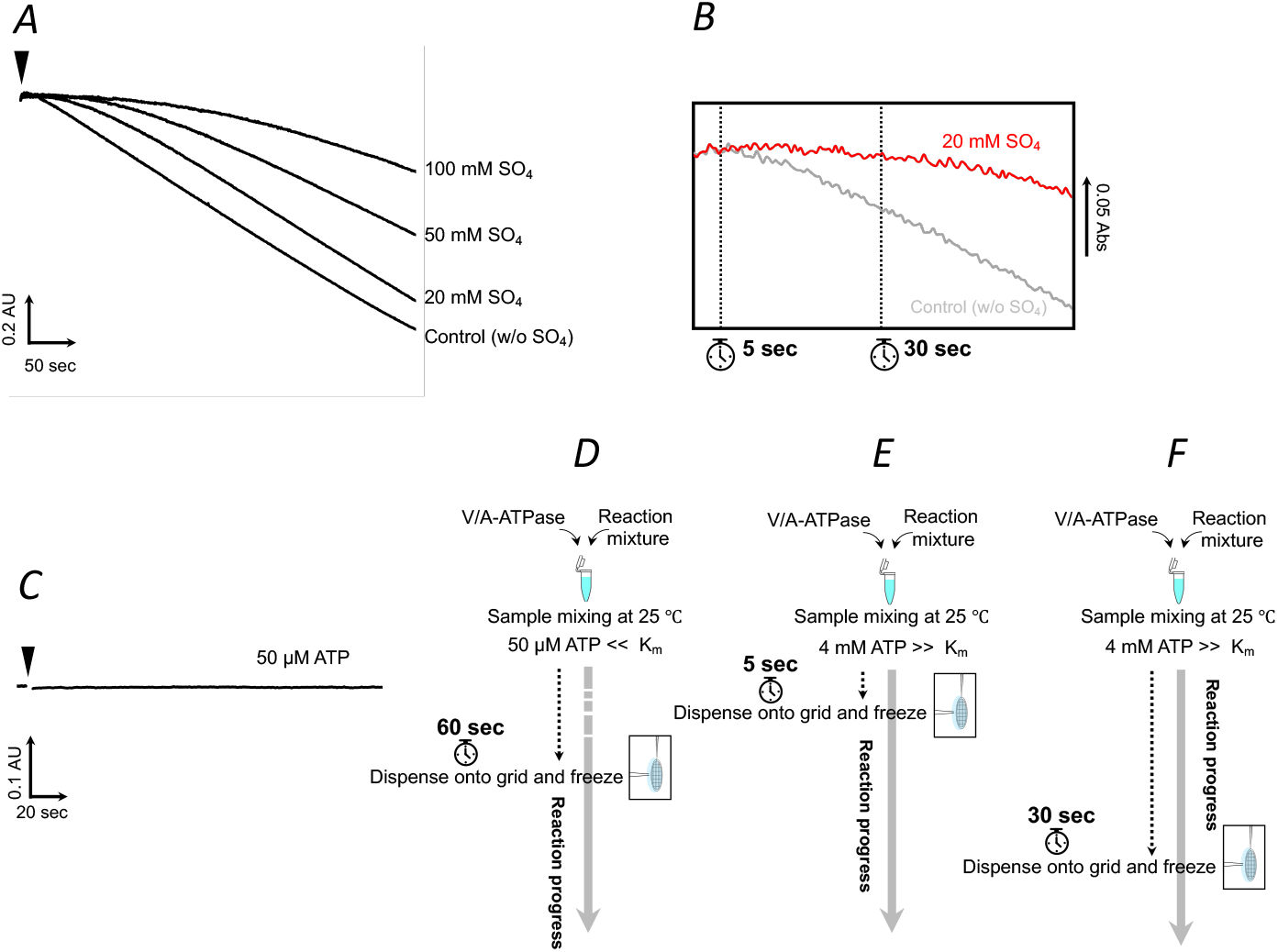
Sulfate extends the duration of the initial intermediates of the V/A-ATPase. **A**, Time-course of ATP hydrolysis catalyzed by the nucleotide-free V/A-ATPase (V_nucfree_) under saturating ATP (a final concentration of 4mM ATP) condition in the absence (Control) or presence of sulfate at various concentrations as indicated. The ATPase activity of V_nucfree_ in the presence of 20 mM sulfate was 18 s^−1^, approximately equal to that of the control (17 s^−1^). ATPase activity of the enzyme in the presence of 50 mM or 100 mM sulfate was 14 or 12 s^−1^, respectively, where the ATPase activity was partially inhibited. The reaction was initiated by adding 20 pmol enzymes (shown by an arrowhead) to 2 mL of the assay mixture. **B**, Magnified view of the initial lag phase in **A. C**, Time-course of ATP hydrolysis catalyzed by V_nucfree_ under much lower ATP concentration than *K*_m_ value (a final concentration of 50 μM ATP) in the presence of sulfate (a final concentration of 20 mM SO_4_). **D-F**, the experimental setup for EM grid preparation. **D**, 20 μM of ATP was added to the reaction mixture containing enzymes and incubated for 60 s at 25 °C before plunge freezing. **E, F**, 4mM of ATP was added to the reaction mixture containing enzymes and incubated for 5 sec (**E**) or 30 sed (**F**) at 25 °C before plunge freezing.

### Structure of V_semi1ATP_ following a 120° step by binding of ATP on the first catalytic site

Nanodiscs reconstituted with V_nucfree_ at a final concentration of 4 μM were mixed with ATP at final concentrations of 50 μM and 20 mM sulfate in the presence of an ATP regeneration system, and the mixture was then incubated at 25 °C for 60 s (Figure 2D). The incubated reaction mixture was then applied to an EM grid and plunge-frozen in liquid ethane. The frozen enzymes were imaged by cryo-EM, followed by image processing using standard procedures (Figure 3-figure supplement 1). The 1,452 K particles were classified into three conformations, termed “state1” at 2.7 Å, “state2” at 3.1 Å, and “state3” at 3.2 Å, which were related by a rotation of DF subunit as shown in a previous study [23, 24, 33]. To improve the local resolution of the V_1_ domain, the density of the membrane-embedded region of state1 was subtracted using the mask covering the V_1_ domain with the part of two stalk regions of the EG. Following refinement of the subtracted particles, we specified the mask range including the AB_semi_ and A_open_ domains for a masked 3D classification procedure, thereby enabling the generation of cryo-EM maps of two different states at 2.8 and 3.2 Å resolution, respectively (Figure 3-figure supplement 1).

The cryo-EM map at 2.8 Å resolution exhibited the asymmetrical structure of the A_3_B_3_ hexamer, as previously shown [23, 24], and had no nucleotide binding to AB_open_. In contrast, spherical densities corresponding to sulfate ions were observed at the catalytic sites on both AB_semi_ and AB_closed_ (Figure 3A). We termed this structure V_2SO4_. For another cryo-EM map at 3.2 Å resolution, the densities of a sulfate ion and ATP were observed in AB_closed_ and AB_semi_, respectively (Figure 3C). However, there was no density at the catalytic site of AB_open_, where the first ATP likely binds as the first step of the reaction. The structure containing ATP at the AB_semi_ is termed V_semi1ATP_.

It is unlikely that ATP binds directly to the catalytic site of AB_semi_ because of its closed structure. Therefore, ATP at the catalytic site of AB_semi_ is most likely derived from ATP bound to AB_open_. This indicates that V_semi1ATP_ is in the state after 120° rotation of the axis, following only one ATP binding to the catalytic site on AB_open_ of V_2SO4_. This expectation was supported by the results of the time-resolved experiment for V/A-ATPase under saturating ATP conditions, as described in the next chapter.

**Figure 3.**
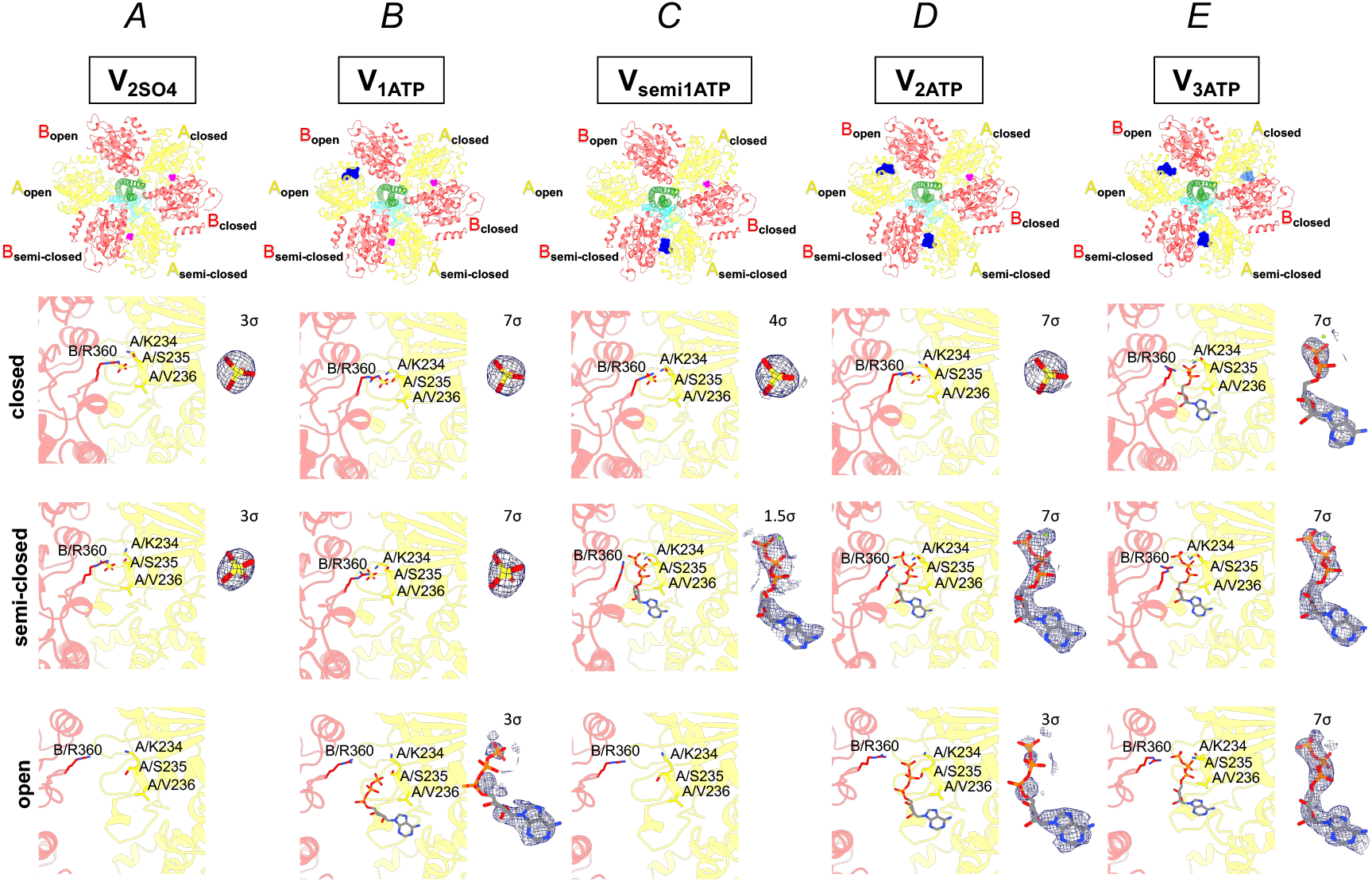
Distinct nucleotide occupancy statuses between intermediates of each condition. Cross sections of each nucleotide-binding site of V_2SO4_ **(A)**, V_1ATP_, (**B**), V_semi1ATP_ (**C**), V_2ATP_ **(D)**, and V_3ATP_ **(E)** viewed from the top are indicated in the top. ATP, ADP, and sulfate are colored blue, light blue, and magenta, respectively. Subunits are colored and labeled with text in the same color. Nucleotide-binding sites and density for nucleotides or sulfates bound at nucleotide-binding sites are shown at bottom. Each contour level for density shown as the blue mesh is indicated. Residues of B/R360, A/K234, A/S235, and A/V236 that interact with nucleotide or sulfate are shown as sticks. The map reveals relative density to nucleotides or sulfates on catalytic sites of each condition and no apparent extra density on the catalytic site of AB_open_ on V_2SO4_ and V_semi1ATP_.

### Structures of the V/A-ATPase incubated with saturating ATP for 5 s

The profile of ATPase activity of V/A-ATPase at 4 mM ATP indicates a lag time corresponding to the slow activation process of the initial inactivated intermediates (Figure 2A). The activation process of V/A-ATPase was not completed within approximately 60 s, which is suitable for trapping intermediates at the indicated time. Therefore, V/A-ATPase was added to the reaction solution and incubated for 5 and 30 s, respectively (Figure 2E, F). Each sample was applied to an EM grid and imaged by cryo-EM, followed by single-particle analysis (Figure 3-figure supplement 2, 3). Particle images (86 K) of state1 were selected after 3D classification in the dataset resulting from the 5-sec reaction mixture. Two classes were classified from the chosen particle images by masked 3D classification focused on the AB_semi_ and A_open_ domains (Figure 3-figure supplement 2). The structure referred to as V_1ATP_ reconstituted from 54 K particles showed ATP binding to the catalytic site on AB_open_ and sulfate binding to the catalytic sites on both AB_closed_ and AB_semi_ (Figure 3B). In structures reconstituted from 30 K particles, ATP molecules were identified in the catalytic sites on both AB_open_ and AB_semi_, and sulfate was confirmed in the catalytic site on AB_closed_ (Figure 3D). This structure was termed V_2ATP_. The V_1ATP_ is an intermediate structure formed immediately after ATP binding to the AB_open_ of V_2SO4_. V_2ATP_ is an intermediate structure formed immediately after ATP binding to V_semi1ATP_, since the catalytic site on AB_open_ of V_semi1ATP_ is immediately occupied by ATP under saturating ATP conditions. These results suggest that ∼35% of the particles of V_1ATP_ transition to V_semi1ATP_ by a 120° rotation of the axis, followed by ATP binding to V_semi1ATP,_ resulting in the generation of V_2ATP_.

### Structures of V/A-ATPase incubated with saturated ATP for 30 s

After 3D masked classification with a mask containing AB_semi_, two structures were obtained from the resulting dataset of the 30 s reaction mixture (Figure 3-figure supplement 3). The structure reconstructed from 16 K particles was identical to the V_1ATP_ obtained from the dataset of the 5 s reaction (Figure 3-figure supplement 2, 3). In another structure reconstructed from 19 K particles, ATP or ADP occupied all three catalytic sites. The structure of the V/A-ATPase containing three nucleotides referred as V_3ATP_ was identical to V_3nuc_ obtained under saturating ATP conditions in our previous study [23] (Figure 3E, Figure 3-figure supplement 3). By contrast, V_2ATP_ disappeared under these conditions. These results indicate that all particles of V_2ATP_ change their state to V_3ATP/3nuc_. For this transition, the 120° rotation of the rotor should occur in V_2ATP_, before ATP binding to empty AB_open_, resulting in the generation of V_2nuc_ structure (empty AB_open_, AB_semi_ with ATP, and AB_closed_ with ADP and Pi). However, neither V_2ATP_ nor other intermediates were identified under these conditions. In contrast, V_1ATP_ was still identified in the dataset from the reaction mixture 30 s after reaction initiation, suggesting that the transition from V_1ATP_ to V_semi1ATP_ did not occur immediately. Together, these results suggest that 120° rotation does not occur immediately by the zippering motion of Ab_open_ with ATP in the absence of ATP binding on AB_semi_. This is consistent with the results of a previous study, in which both the zippering motion of AB_open_ with ATP and hydrolysis of ATP in AB_semi_ drive the unzipper motion of AB_closed_ with the release of ADP and P*i*, which are coupled simultaneously with the 120° rotation of the shaft [23].

### Different conformations of AB_semi_ with different ligands

After the refinement with signal subtraction for the membrane domain, different conformations of AB_semi_ were identified from the masked 3D classification procedure (Figure 3-figure supplement 1-3). C_α_ root means square displacement (RMSD) values between semi-closed B subunits indicated a significant structural difference in the C-terminal helix bundle between V_1ATP_ and V_semi1ATP_ when superimposed on the N-terminal β-barrel domain (Supplementary table 1). The C-terminal helix bundle of the B subunit changed to a more open conformation upon ATP binding (Figure 3-figure supplement 4A, black square area). In contrast, there was no significant difference between the semi-closed A subunits (Figure 3-figure supplement 4, top). Considering the difference in ligands between these states, ATP binding induces conformational changes in the C-terminal helix bundle of the semi-closed B subunit.

Furthermore, the contour levels on the maps for ATP binding to AB_semi_ varied between the states (Figure 3). Two maps of V_2ATP_ and V_3ATP_ showed an apparent density for ATP binding to each catalytic site on AB_semi_ when contoured at 7σ (standard deviation). In contrast, the density for ATP binding to AB_semi_ in V_semi1ATP_ had to be lowered to 1.5σ for visualization (Figure 3C, D, E). The atomic displacement (B) factor of ATP binding to AB_semi_ in the V_2ATP_ and V_3ATP_ was in the range of 30–59 and 41–77 Å^2^, respectively, although V_semi1ATP_ showed a higher B factor range of 112–133 Å^2^. The low contour threshold of ATP binding to the AB_semi_ of V_semi1ATP_ is associated with a high B factor, which implies greater flexibility and/or heterogeneity within the nucleotide. These results imply multiple ligand conformations of the nucleotide binding to the catalytic site on AB_semi_ of V_semi1ATP_ in the initial reaction process.

## Discussion

Cryo-EM snapshot analysis of V/A-ATPase indicated that the 120° step of the axis is driven by three reactions occurring simultaneously at the three catalytic sites [23]. This raises questions on how nucleotide-free V/A-ATPase transitions to a steady state by ATP binding to the catalytic sites on the enzyme. Previous studies have suggested the existence of an initial inactivated state of V_1_-ATPase [29]. In this study, we attempted to capture short-lived initial intermediates of V/A-ATPase in the presence of sulfate, which can temporally stabilize the initial intermediate.

In the presence of 50 μM ATP and 20 mM sulfate, structures of both V_2SO4_ and V_semi1ATP_ were obtained, where AB_closed_ was occupied by sulfate. The binding of sulfate to the catalytic site on AB_closed_ is likely to stabilize its closed conformation, hampering the 120° step of the axis by the zippering motion of AB_open_ with ATP. Nevertheless, our results indicated that V_2SO4_ and V_semi1ATP_ finally transformed to V_3ATP_ (ATP/ADP occupies all three AB) by sequential binding of ATP to the catalytic sites (Figure 4). This observation suggests that sulfate binding to the catalytic site on AB_closed_ weakly stabilizes the closed structure of the catalytic site. In contrast, the ADP-inhibited state of the wild-type V/A-ATPase is defective in hydrolyzing ATP, where the unzippering motion of AB_closed_ is hampered by ADP binding to the catalytic site on AB_closed_ [23, 24].

**Figure 4.**
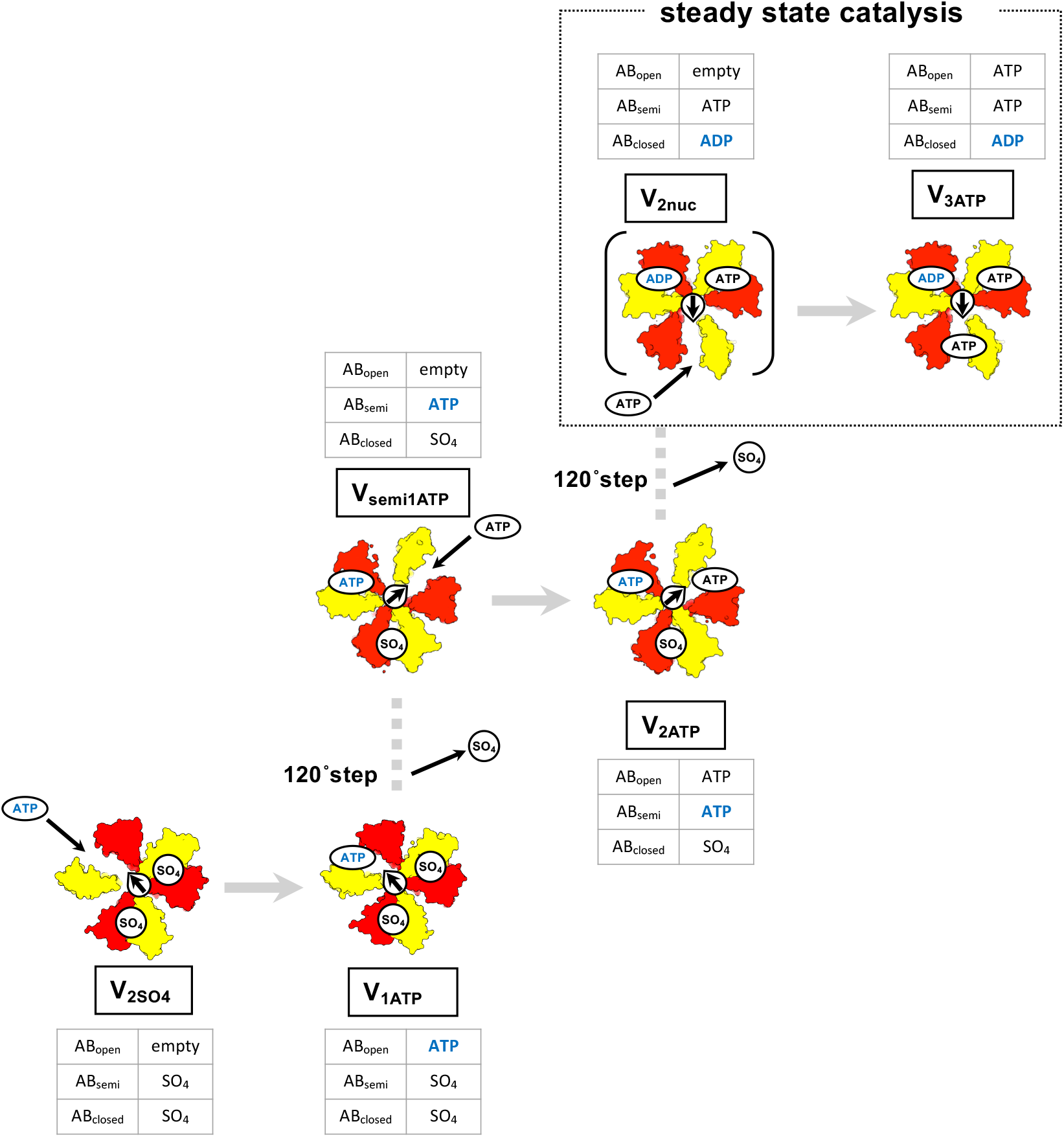
The activation process of the ground state of the V/A-ATPase. Cryo-EM maps for each state are shown as surfaces, colored as in Fig. 1. DF axis is shown as an arrow to clarify the scheme, and corresponding tables for all AB pairs observed in the structures are indicated. In the initial phase, V_2SO4_ is activated by binding ATP colored blue to the AB_open_, then changes its conformation to V_1ATP_ (lower left). ATP binding to the AB_open_ in V_1ATP_ induces the release of sulfate binding to the AB_closed_ with the axis rotation in 120° step (to the middle from lower left). Consequently, V_1ATP_ changes its conformation to V_semi1ATP_. V_semi1ATP_ changes its conformation to V_2ATP_ in succession following ATP binding to the AB_open_ (middle). ATP binding to V_semi1ATP_ is expected to induce the release of remaining sulfate binding to the AB_closed_ with the axis rotation in 120° step (to the right upper from the middle), and finally, V_2ATP_ changes its conformation to V_3ATP_ via V_2nuc_ and enters the continuous catalytic cycle.

Under the waiting conditions for ATP binding ([ATP] < *K*_m_), AB_open_ in the V/A-ATPase adopts an empty form since ATP binding to the catalytic site on AB_open_ is a rate-limiting step and other catalytic reaction(s) are mostly completed while waiting for ATP binding. Consequently, V_semi1ATP_ was obtained under waiting conditions for ATP binding instead of V_1ATP_, whose catalytic site on AB_open_ is occupied by ATP. V_semi1ATP_ is the state following 120° rotation of the rotor in V_1ATP_ (Figure 4, from the middle to the lower left). This suggests that ATP binding to AB_open_ can initiate a 120° step of the axis when an inhibitory ADP does not occupy AB_closed_.

Both V_1ATP_ and V_2ATP_ were identified in the resulting dataset from the reaction solution after 5 s under the ATP-saturating condition ([ATP] > *K*_m_). The catalytic sites on AB_open_ in both V_1ATP_ and V_2ATP_ are filled with ATP because ATP binding to AB_open_ is not a rate-limiting step. Consequently, V_1ATP_ is apparently in the state immediately after ATP binds to V_2SO4_ (Figure 4, lower left). Similarly, V_2ATP_ is in the state immediately after ATP binding to the catalytic site on AB_open_ in V_semi1ATP_ (Figure 4, middle). V_2ATP_ disappeared in the resulting dataset from the reaction solution after 30 s, while V_1ATP_ was still identified (Figure 3-figure supplement 3), indicating that V_2ATP_ is a short-lived intermediate compared to V_1ATP_. This observation implies that ATPase activity lagged because of a delayed transition from V_1ATP_ to V_2ATP_ via V_semi1ATP_ in the initial reaction. Instead, V_3ATP_ would be identified in the same dataset. The V_3ATP_ is likely derived from V_2nuc_ where AB_open_ is empty (Figure 4, to upper right from middle, [23]). Once all V_2ATP_ change to V_2nuc_, the steady state catalysis continues by the tri-site mechanism [Figure 4, upper].

Assuming that the structural transition from V_1ATP_ to V_semi1ATP_ is a rate-limiting step in the activation process, V_1ATP_ is likely to be identified even at 50 μM ATP. However, approximately 95% of the particles were classified as V_2SO4_, and V_1ATP_ was not identified under 50 μM ATP. The particles of the V_1_ domain are classified into different intermediates according to the difference in the AB_semi_ structure due to the bound ligand (Figure 3-figure supplement 4). Therefore, it is possible that the V_1ATP_ particles were not distinguished from the V_2SO4_ particles, and the V_1ATP_ structure was not identified.

Our structural snapshots revealed that the three AB conformations, referred to as “open,” “closed,” and “semi-closed” form, for each three catalytic AB pair of the V/A-ATPase was maintained during the activation process of the ground state enzyme, regardless of nucleotide binding status. A recent structural study of bacterial F_o_F_1_ using cryo-EM single particle analysis revealed open conformations of all three catalytic β subunits in the nucleotide-depleted enzyme [25]. This structure significantly differs from the canonical F_1_ structure, which contains two closed β subunits and one open β subunit. Furthermore, the first ATP under uni-site conditions was hydrolyzed at β_TP_, followed by a structural transition of open β_TP_ to closed β_DP_. The F_o_F_1_ after uni-site catalysis adopts a structure with two open β and one closed β subunit. The structural plasticity of the F_1_ domain is in contrast to the structural rigidity of the V_1_ domain, which maintains the same asymmetric structure independent of nucleotide occupancy at the catalytic sites.

In summary, we identified intermediate structures that appear during the transition from the initial state to the steady state, which cannot be obtained by canonical structural analysis using cryo-EM or X-ray crystallography. Due to the limitation of current experimental techniques, a detailed knowledge of conformational changes in molecular motor is missing. In addition to the analysis of the steady-state function of the V/A-ATPase, elucidating the activation process of the enzyme will pave the way for a complete understanding of the molecular mechanism of the rotational molecular motor.

## Methods

### Preparation of V/A-ATPase for Biochemical Assays and Cryo-EM Imaging

The V/A-ATPase containing His3 tags on the C-terminus of each c subunit and the TSSA mutation (S232A and T235S) on the A subunit were isolated from *T. thermophilus* membranes as previously described [14], with certain modifications. The enzyme solubilized from the membranes with 10% Triton X-100 was purified by Ni^2+^-NTA affinity with 0.03% dodecyl-*β-D*-maltoside (DDM). For bound nucleotide removal, the eluted fractions containing V/A-ATPase were dialyzed against 200 mM sodium phosphate (pH 8.0), 10 mM EDTA, and 0.03% DDM overnight at 25 °C with three buffer changes, followed by dialysis against 20 mM Tris-Cl (pH 8.0), 1 mM EDTA, and 0.03% DDM (TE buffer) before anion exchange chromatography using a 6 mL Resource Q column (GE Healthcare). Nucleotide-free *Tth* V/A-ATPase (V_nucfree_) was eluted with a linear NaCl gradient using TE buffer (0–500 mM NaCl, 0.03% DDM). The eluted fractions containing V_nucfree_ were concentrated to ∼10 mg/mL using Amicon 100k molecular weight cut-off filters (Millipore). For nanodisc incorporation, 1,2-Dimyristoyl-sn-glycero3-phosphorylcholine (DMPC, Avanti) was used to form lipid bilayers in reconstruction, as previously described [23, 33]. Purified V_nucfree_ solubilized in 0.03% n-DDM was mixed with the lipid stock and membrane scaffold protein MSP1E3D1 (Sigma) at a specific molar ratio of enzyme: MSP: DMPC lipid = 1:4:520 and incubated on ice for 0.5 h. Then, 200 μL of Bio Beads SM-2 equilibrated with a wash buffer (20 mM Tris-HCl, pH 8.0, 150 mM NaCl) was added to the 500 μL mixture. After 2 h of incubation at 4 °C with gentle stirring, an additional 300 μL of Bio Beads was added, and the mixture was incubated overnight at 4 °C to form the nanodiscs. The supernatant of the mixture containing V_nucfree_ reconstituted into nanodiscs was loaded onto a Superdex 200 Increase 10/300 column equilibrated with the wash buffer. The peak fractions were collected, analyzed by sodium dodecyl sulfate-polyacrylamide gel electrophoresis, and concentrated to 4 mg/mL. The prepared ND-*Tth* V/A-ATPase was immediately used for biochemical assays or cryo-grid preparation, as V_nucfree_ aggregates within a few days.

### Biochemical assay

ATPase activity was measured at 25 °C using an enzyme-coupled ATP-regenerating system as described previously [14, 23]. The reaction mixture contained 50 mM Tris-HCl (pH 8.0), 100 mM KCl, different concentrations of ATP-Mg, 2.5 mM phosphoenolpyruvate (PEP), 50 μg/mL pyruvate kinase (PK), 50 μg/mL lactate dehydrogenase, and 0.2 mM NADH in a final volume of 2 mL. The reaction was initiated by adding 20 pmol of enzymes to 2 mL of the assay mixture with or without sulfate, and the rate of ATP hydrolysis was monitored, as the rate of oxidation of NADH was determined by the absorbance decrease at 340 nm.

### Cryo-EM imaging

Sample vitrification was performed using a semi-automated device (Vitrobot, FEI). The reaction basal buffer (RB buffer) containing 50 mM Tris-Cl (pH 8.0), 100 mM KCl, 2 mM MgCl_2_, 5 mM PEP, 100 μg/mL PK, and 20 mM ammonium sulfate was used under different reaction conditions. For V_nucfree_ with a low concentration of ATP, 4 μM of the enzyme was mixed with the same volume of ×2 RB buffer containing 100 μM ATP-Mg. Then the mixtures were incubated for 60 sec at 25 °C; then 2.4 μL of sample mixture was transferred to a glow-discharged grid (Quantifoil R1.2/1.3, 300 Mesh, molybdenum) (Figure 2D). The grid was then automatically blotted once from both sides with filter paper for a 6 s blot time. The grid was then plunged into liquid ethane with no delay. For V_nucfree_ with saturated ATP, 4 μM of the enzyme was mixed with the same volume of ×2 RB buffer containing 8 mM ATP-Mg. The mixtures were incubated for 5 or 30 s at 25 °C to resolve intermediates during the initial phase, followed by blotting and vitrification (Figure 2E, F).

For V_nucfree_, under the waiting conditions for ATP binding ([ATP] < *K*_m_), cryo-EM movie collection was performed with a CRYOARM 300 (JEOL) operating at an accelerating voltage of 300 keV and equipped with a K3 (Gatan) direct electron detector in electron counting mode (CDS, [34]), using the data collection software serialEM [35]. The pixel size was 0.81 Å/pix (×60,000), and a total dose of 50.0 e^−^ Å^−2^ (1.0 e^−^ Å^−2^ per frame) with a total 1.5 sec exposure time (50 frames) with a defocus range of −1.2 to −2.4 μm. The image processing steps for the reaction conditions are summarized in Figure 3-figure supplement 1.

For time-resolved data in V_nucfree_ under saturating ATP conditions, cryo-EM imaging was performed using a Titan Krios (FEI/Thermo Fisher) operating at an acceleration voltage of 300 kV and equipped with a direct K3 (Gatan) electron detector in electron counting mode (CDS, [34]). Data collection was carried out using SerialEM software [35] at a calibrated magnification of 0.83 Å pixel^−1^ (×105,000) and a total dose of 50.0 e^−^ Å^−2^ (or 1.0 e− Å^−2^ per frame) (where e^−^ specifies electrons) with a total 5 sec exposure time. The defocus range was −0.8 to −1.8 μm. Data were collected from 48 video frames. Image processing steps for 5 and 30 s time points are summarized in Figure 3-figure supplements 2 and 3, respectively.

### Image processing

Image processing steps for each reaction condition are summarized in Figure 3-figure supplement 1-3. Collected data were processed with RELION [36], and Cryosparc [37], motion-corrected with the MotionCor2 algorithm [38]; Initial CTF parameters from each dose-weighted image were determined by using CTFFIND4 [39]. The details were identical to those described in our previous paper [23]. For three datasets, several rounds of two-dimensional classification with 5x binned particles using C1 symmetry were performed to select intact V/A-ATPase particles. Then they were applied to 3D classification to separate three states (state1-3), which were related by a rotation of the DF subunit. Each selected particle was re-extracted at full pixel size and applied to 3D autorefinements, followed by CTF refinement and Bayesian polishing. The final maps of the holo-complex were generated using another round of CTF refinement and 3D autorefinements using soft masks. The membrane domain was visible but less clear than the hydrophilic V_1_ domain in a holo-enzyme map. This seemed to be due to the structural flexibility resulting from the movement of the membrane domain relative to V_1_ [23, 24, 33]. To improve the local resolution of the V_1_ domain, the density of the membrane-embedded region of state 1 was subtracted by using the mask covering the V_1_ and C-terminal domain of EG subunits (V_1_EG). The refinements provided the density maps for V_1_EG under each condition at 2.4–3.0 Å resolution (Figure 3-figure supplement 1-3).

For V_1_EG under the condition of a low ATP concentration dataset, refined particles (216,617 particles in total) were applied to additional rounds of 3D no-alignment classification (regularization parameter *T* = 20, the number of classes *k* = 6) with a mask containing the semi-closed AB and the open A subunit to identify substrates after conventional 3D classification. Among these classes, 145,880 particles belonging to V_2SO4_ and 39,101 particles belonging to V_semi1ATP_ were selected for further refinement.

Similar procedures were used to process V_1_EG with a saturating ATP dataset. For the time-resolved dataset in V_1_EG at 5-sec time point with saturating ATP, refined particles (86,828 particles in total) were applied to additional rounds of 3D no-alignment classification (regularization parameter *T* = 20, the number of classes *k* = 5) with a mask containing the AB_semi_ and the open A subunit to identify substrates after conventional 3D classification. Among these classes, 54,047 particles belonging to V_1ATP_ and 30,284 particles belonging to V_2ATP_ were selected for further refinement.

For the time-resolved dataset in V_1_EG at the 30 s time point with saturating ATP, refined particles (46,901 particles in total) were applied to additional rounds of 3D no-alignment classification (regularization parameter *T* = 32, the number of classes *k* = 4) with a mask containing the AB_semi_ and the open A subunit to identify substates after conventional 3D classification. Among these classes, 19,234 particles of V_3ATP_ and 15,867 particles of V_1ATP_ were selected for further refinement.

The soft masks of V_1_EG for different substrates were further applied to the continued 3D autorefinements, resulting in the overall resolution reported in Supplementary table 2 after post-processing, including soft masking and B-factor sharpening. Alternative post-sharpening was performed using DeepEMhancer [40]. The resolution was based on the gold-standard Fourier shell correlation criterion (0.142). The local resolution was calculated using ResMap [41]. Two maps of V_1ATP_ identified from different datasets were nearly identical; hence, a higher resolution map of V_1ATP_ from the 5 s time point dataset was used for further analysis.

### Model building, refinement, and validation

Maps from DeepEMhancer were used for model building, refinement, and subsequent structural interpretation. The V_1_EG model of ND-*Tth* V/A-ATPase was adopted from the 3.1 Å V_1_EG structure (PDB accession no. 7VAL) as the initial model [23]. With 2.5-3.1 Å V_1_EG density maps, we could model side chains and local geometry to achieve higher accuracy (Figure 3-figure supplement 5). The initial model was docked into the cryo-EM map using UCSF ChimeraX [42], and sulfate ions were built into COOT [43], followed by several rounds of real-space refinement using Phenix [44, 45]. The initial model manually corrected residue by residue in COOT [43] regarding side-chain conformations. The iterative process using COOT [43] and ISOLDE [46] was performed for several rounds to correct the remaining errors until the model agreed well with the geometry (Supplementary table 2). Cross-validation was carried out by comprehensive validation (cryo-EM) in Phenix [44, 45], and the map to model FSC curves was calculated (Figure 3-figure supplement 5B). RMSD values between the atomic models and structural figures were calculated using the UCSF chimeraX [42].

## Acknowledgements

We are grateful to all the members of the Yokoyama Lab and Mitsuoka Lab for their continuous support and technical assistance. We also thank Dr. T Nishizawa and Dr. F Makino affiliated with University of Tokyo (present affiliation; Yokohama City University) and JEOL Ltd, respectively, for helping with the electron microscopy data collection.

## Author contributions

Author contributions are as follows; AN, KY and JK analyzed the data and contributed to the preparation of the samples. KM provided technical support and conceptual advice. KY and AN designed and supervised the experiments and wrote the manuscript. All authors discussed the results and commented on the manuscript.

## Data and material availability

Cryo-EM density maps and corresponding atomic models have been deposited in the Electron Microscopy Data Bank (EMDB) and Protein Data Bank (PDB), respectively, with the following accession numbers (EMDB, PDB): V_1ATP_ (EMD-34362, 8GXU), V_2ATP_ (EMD-34363, 8GXW), V_3ATP_ (EMD-34364, 8GXX), V_2SO4_ (EMD-34365, 8GXY), V_semi1ATP_

(EMD-34366, 8GXZ). The data that support the findings of this studies are available from the corresponding author upon reasonable request.

## References

1. Kuhlbrandt, W. Structure and Mechanisms of F-Type ATP Synthases. Annu Rev Biochem 88, 515–549 (2019)

2. Forgac, M. Vacuolar ATPases: rotary proton pumps in physiology and pathophysiology. Nat Rev Mol Cell Biol 8, 917–929 (2007)

3. Yoshida, M., Muneyuki, E. & Hisabori, T. ATP synthase—a marvellous rotary engine of the cell. Nat. Rev. Mol. Cell Biol. 2, 669–677 (2001)

4. Boyer, P. D. The ATP synthase--a splendid molecular machine. Annu Rev Biochem 66, 717–749 (1997)

5. Kühlbrandt, W. & Davies, K. M. Rotary ATPases: A New Twist to an Ancient Machine. Trends. Biochem. Sci. 41, 106–116 (2016)

6. Yokoyama, K. & Imamura, H. Rotation, structure, and classification of prokaryotic V-ATPase. J Bioenerg Biomembr 37, 405–410 (2005)

7. Yokoyama, K., Oshima, T., Yoshida, M. Thermus thermophilus membrane-associated ATPase. Indication of a eubacterial V-type ATPase J. Biol. Chem 265, 21946–21950 (1990)

8. Imamura, H., Nakano, M., Noji, H., Muneyuki, E., Ohkuma, S., Yoshida, M., Yokoyama, K. Evidence for rotation of V_1_-ATPase. Proc. Natl. Acad. Sci. USA 100, 2312–2315 (2003)

9. Muench, S. P., Trinick, J., Harrison, M. A. Structural divergence of the rotary ATPases. Q. Rev. Biophys. 44, 311–356 (2011)

10. Kane, P. M. Disassembly and reassembly of the yeast vacuolar H(+)-ATPase in vivo. J. Biol. Chem. 270:17025–17032. (1995)

11. Kishikawa, J., Nakanishi, A., Furuike, S., Tamakoshi, M., Yokoyama, K. Molecular basis of ADP inhibition of vacuolar (V)-type ATPase/synthase J. Biol. Chem. 289(1):403–412. doi: 10.1074/jbc.M113.523498. (2014)

12. Stewart, A. G., Sobti, M., Harvey, R. P., Stock, D. Rotary ATPases: models, machine elements and technical specifications Bioarchitecture. 3:2–12. doi: 10.4161/bioa.23301. (2013)

13. Mazhab-Jafari, M. T. & Rubinstein, J. L. Cryo-EM studies of the structure and dynamics of vacuolar-type ATPases. Sci Adv 2, e1600725 (2016)

14. Nakano, M., Imamura, H., Toei, M., Tamakoshi, M., Yoshida, M., Yokoyama, K. ATP hydrolysis and synthesis of a rotary motor V-ATPase from Thermus thermophilus. J. Biol. Chem. 283, 20789–20796 (2008)

15. Furuike, S., Nakano, M., Adachi, K., Noji, H., Kinoshita, K. Jr., Yokoyama, K. Resolving stepping rotation in Thermus thermophilus H+ -ATPase/synthase with an essentially dragfree probe. Nat. Commun. 2, 233 (2011)

16. Iwata, M., Imamura, H., Stambouli, E., Ikeda, C., Tamakoshi, M., Nagata, K., Makyio, H., Hankamer, B., Barber, J., Yoshida, M., Yokoyama, K., Iwata, S. Crystal structure of a central stalk subunit C and reversible association/dissociation of vacuole-type ATPase. Proc. Natl. Acad. Sci. USA 101, 59–64 (2004)

17. Makyio, H., Iino, R., Ikeda, C., Imamura, H., Tamakoshi, M., Iwata, M., Stock, D., Bernal, R. A., Carpenter, E. P., Yoshida, M., Yokoyama, K., Iwata, So. Structure of a central stalk subunit F of prokaryotic V-type ATPase/synthase from Thermus thermophilus. EMBO J. 24, 3974–3983 (2005)

18. Lee, L. K., Stewart, A. G., Donohoe, M., Bernal, R. A., Stock, D. The structure of the peripheral stalk of Thermus thermophilus H+ -ATPase/synthase. Nat. Struct. Mol. Biol. 17, 373–378 (2010)

19. Murata, T., Yamato, I., Kakinuma, Y., Leslie, A. G., Walker, J. E. Structure of the Rotor of the V-Type Na+ -ATPase from Enterococcus hirae. Science 308, 654–659 (2005)

20. Srinivasan, S., Vyas, N. K., Baker M. L., Quiocho, F. A. Crystal structure of the cytoplasmic N-terminal domain of subunit I, a homolog of subunit a, of V-ATPase J. Mol. Biol. 412(1)14–21 doi: 10.1016/j.jmb.2011.07.014 (2011)

21. Maher, M.J., Akimoto, S., Iwata, M., Nagata, K., Hori, Y., Yoshida, M., Yokoyama, S., Iwata, S., Yokoyama, K. Crystal structure of A3B3 complex of V-ATPase from Thermus thermophilus. EMBO J. 28, 3771–3779 (2009)

22. Nagamatsu, Y., Takeda, K., Kuranaga, T., Numoto, N., Miki, K. Origin of Asymmetry at the Intersubunit Interfaces of V_1_-ATPase from Thermus thermophilus. J. Mol. Biol. 425, 2699–2708 (2013)

23. Kishikawa, J., Nakanishi, A., Nakano, A., Saeki, S., Furuta, A., Kato, T., Mitsuoka, K., Yokoyama, K. Structural snapshots of V/A-ATPase reveal the rotary catalytic mechanism of rotary ATPases Nature Commun. 13(1):1213. doi: 10.1038/s41467-022-28832-5 (2022)

24. Nakanishi, A., Kishikawa, J. I., Tamakoshi, M., Mitsuoka, K. Yokoyama, K. Cryo EM structure of intact rotary H(+)-ATPase/synthase from Thermus thermophilus. Nat. Commun. 9, 89 (2018)

25. Nakano, A., Kishikawa, J. I., Nakanishi, A., Mitsuoka, K., Yokoyama, K. Structural basis of unisite catalysis of bacterial F_o_F_1_-ATPase PNAS Nexus, pgac116, doi: 10.1093/pnasnexus/pgac116 (2022)

26. Boyer, P. D. Catalytic site forms and controls in ATP synthase catalysis. Biochim. Biophys. Acta 1458, 252–262 (2000)

27. Löbau, S., Weber, J., Senior, A. E. Nucleotide occupancy of F_1_-ATPase catalytic sites under crystallization conditions. FEBS Lett. 404, 15–18 (1997)

28. Watanabe, R., Noji, H. Timing of inorganic phosphate release modulates the catalytic activity of ATP-driven rotary motor protein Nat. Commun. 5:3486, doi: 10.1038/ncomms4486 (2014)

29. Yokoyama, K., E., Muneyuki., Amano, T., Mizutani, S., Yoshida. M., Ishida, M., Ohkuma, S. V-ATPase of Thermus thermophilus is inactivated during ATP hydrolysis but can synthesize ATP J. Biol. Chem. 273(32):20504–20510. doi: 10.1074/jbc.273.32.20504 (1998)

30. Frank, J. Time-resolved cryo-electron microscopy: recent progress. J. Struct Biol. 200, 303–306 (2017)

31. N. Yoder., F. Jalali-Yazdi., S. Noreng., A. Houser., I. Baconguis., E. Gouaux. Light-coupled cryo-plunger for time-resolved cryo-EM 212(3):107624 J. Struct Biol doi: 10.1016/j.jsb.2020.107624 (2020)

32. Dandey, V. P., Budell, W. C., Wei, H., Bobe, D., Maruthi, K., Kopylov, M., Eng, E. T., Kahn, P. A., Hinshaw, J. E., Kundu, N., Nimigean, C. M., Fan, C., Sukomon, N., Darst, S. A., Saecker, R. M., Chen, J., Malone, B., Potter, C. S., Carragher, B. Time-resolved cryo-EM using Spotiton Nature Methods (9):897–900 doi: 10.1038/s41592-020-0925-6 (2020)

33. Kishikawa, J. I., Nakanishi, A., Furuta, A., Kato, T., Namba, K., Tamakoshi, M., Mitsuoka, K., Yokoyama, K. Mechanical inhibition of isolated V(_o_) from V/A-ATPase for proton conductance. Elife 9, e56862 (2020)

34. Sun, M., Azumaya, C. M., Tse, E., Bulkley, D. P., Harrington, M. B., Gilbert, G., Frost, A., Southeworthe, D., Verba, K. A., Chen, Y., Agard, D. A. Practical considerations for using K3 cameras in CDS mode for high-resolution and high-throughput single particle cryo-EM J. Struct. Biol. 213(3):107745 doi: 10.1016/j.jsb.2021.107745 (2021)

35. Mastronarde, D. N. Automated electron microscope tomography using robust prediction of specimen movements. J. Struct. Biol. 152, 36–51 (2005)

36. Scheres, S. H. RELION: implementation of a Bayesian approach to cryo-EM structure determination. J. Struct. Biol. 180, 519–530 (2012)

37. Punjani, A., Rubinstein, J. L., Fleet, D. J. & Brubaker, M. A. cryoSPARC: algorithms for rapid unsupervised cryo-EM structure determination. Nat. Methods 14, 290–296 (2017)

38. Zheng, S. Q., Palovcak, E., Armache, J-P., Verba, K. A., Cheng, Y., Agard, D. A. MotionCor2: anisotropic correction of beam-induced motion for improved cryoelectron microscopy. Nat. Methods 14, 331–332 (2017)

39. Rohou, A. & Grigorieff, N. CTFFIND4: fast and accurate defocus estimation from electron micrographs. J. Struct. Biol. 192, 216–221 (2015)

40. Sanchez-Garcia, R., Gomez-Blanco, J., Cuervo, A., Carazo, J. M., Sorzano, C. O S., Vargas, J. DeepEMhancer: a deep learning solution for cryo-EM volume post-processing. Commun. Biol. 4, 874 (2020)

41. Kucukelbir, A., Sigworthe, F. J., Tagare, H. D. Quantifying the local resolution of cryo-EM density maps. Nat. Methods. 11, 63–65 (2014)

42. Pettersen, E. F., Goddard, T. D., Huang, C C., Meng, E. C., Couch, G. S., Croll, T. I., Morris, J. H., Ferrin, T. E. UCSF ChimeraX: structure visualization for researchers, educators, and developers. Protein Sci. 30, 70–82 (2021)

43. Emsley, P., Lohkamp, B., Scott, W. G., Cowtan, K. Features and development of Coot. Acta Crystallogr. D. Biol. Crystallogr. 66, 486–501 (2010)

44. Afonine, P. V., Poon, B. K., Read, R. J., Sobolev, O. V., Terwilliger, T. C., Urzhumtsev, A., Adams, P. D. Real-space refinement in PHENIX for cryo-EM and crystallography Acta Crystallogr. D Struct. Biol., 74, 531–544 (2018)

45. Liebschner, D., Afonine, P. V., Baker, M. L., Bunkóczi, G., Chen, V. B., Croll, T. I., Hintze, B., Hung, L. W., Jain, S., McCoy, A. J., Moriarty, N. W., Oeffner, R. D., Poon, B. K., Prisant, M. G., Read, R. J., Richardson, J. S., Richardson, D. C., Sammito, M. D., Sobolev, O. V., Stockwell, D. H., Terwilliger, T. C., Urzhumtsev, A. G., Videau, L. L., Williams, C. J., Adams, P. D. Macromolecular structure determination using X-rays, neutrons and electrons: recent developments in Phenix. Acta Crystallogr. D Struct. Biol. 75, 861–877 (2019).

46. Croll, T. I. ISOLDE: a physically realistic environment for model building into low-resolution electron-density maps. Acta Crystallogr. D. Struct. Biol. 74, 519–530 (2018).

