## Supplemental Files for "Structural transition of the ground-state structure to steady-state structures by sequential binding of ATP to V/A-ATPase"

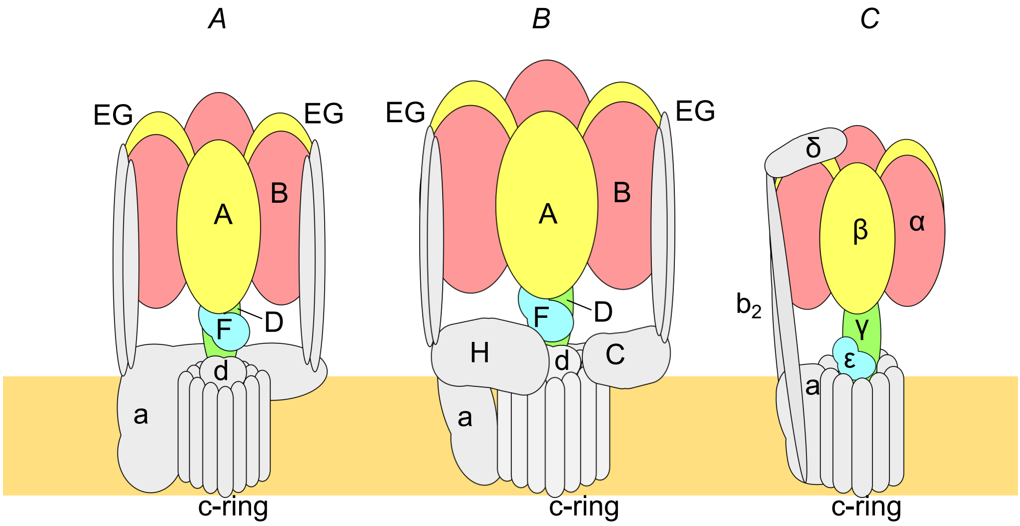

**Figure 1-figure supplement 1. Schematic representation of rotary ATPases.**
Schematic of the subunit composition of different types of rotary ATPases: prokaryotic V/A-ATPase (**A**), eukaryotic V-ATPase (**B**), and prokaryotic F-ATPase (**C**). The hydrophilic $V_1/F_1$ domain is represented by various colors, and the hydrophobic $V_o/F_o$ domain is colored in gray.

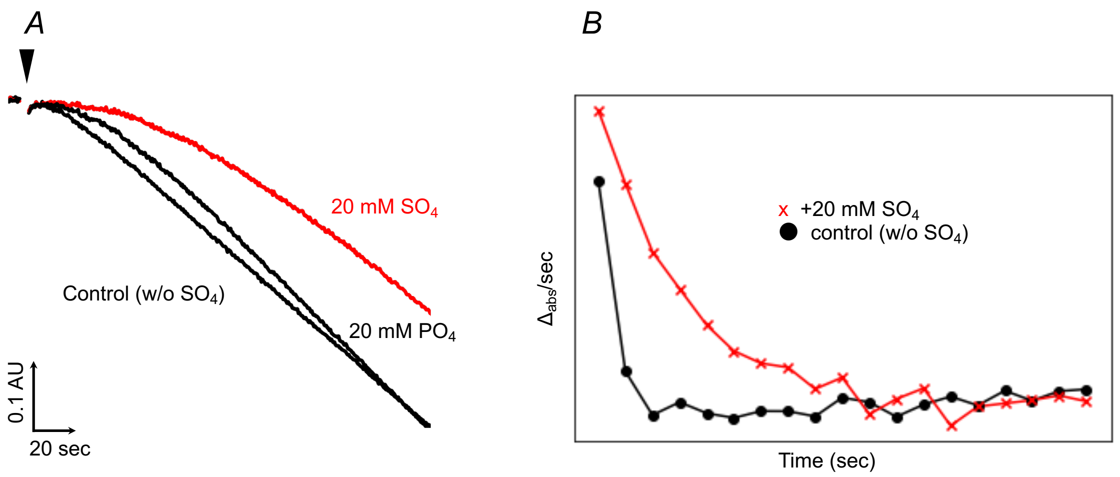

**Figure 2-figure supplement 1. Sulfate effect on ATP hydrolysis by the V/A-ATPase.** **A**, Time-course of ATP hydrolysis catalyzed by the nucleotide-free V/A-ATPase ($V_{nucfree}$) under saturating ATP (a final concentration of 4mM ATP) condition in the absence (Control) or presence of sulfate (20 mM $SO_4$) as indicated. There were almost no changes in ATPase activity due to adding phosphate (20 mM $PO_4$). **B**, Plots show the differential of the ATPase activity per unit time (20 s). It takes even more time to reach steady state in the presence of sulfate (red) in the absence of sulfate (black).

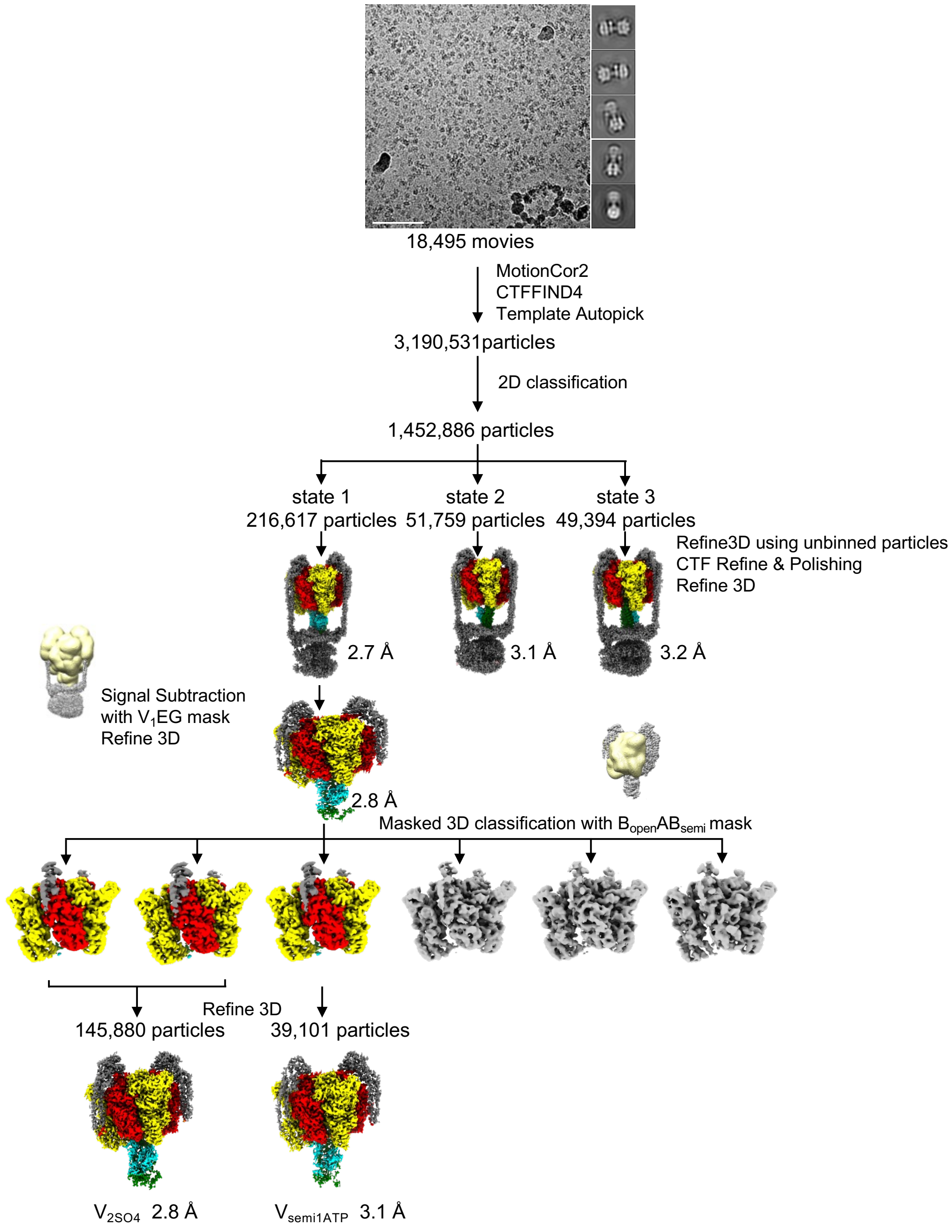

**Figure 3-figure supplement 1. Cryo-EM processing workflow of the V/A-ATPase activated by 20 μM ATP for 60 s.** A processing pipeline for 3D reconstruction of the nucleotide-free V/A-ATPase ($V_{nucfree}$) under much lower ATP concentration than $K_m$ value (a final concentration of 50 μM) with 20 mM sulfate. Example micrograph, 2D class average images (top, scale bar; 100 nm), and cryo-EM data analysis workflow (bottom) are shown. Masked 3D classification allowed us to obtain two other maps with improved quality of the $V_1$EG region at 2.8 Å ($V_{2SO4}$) and 3.1 Å ($V_{semi1ATP}$), respectively.

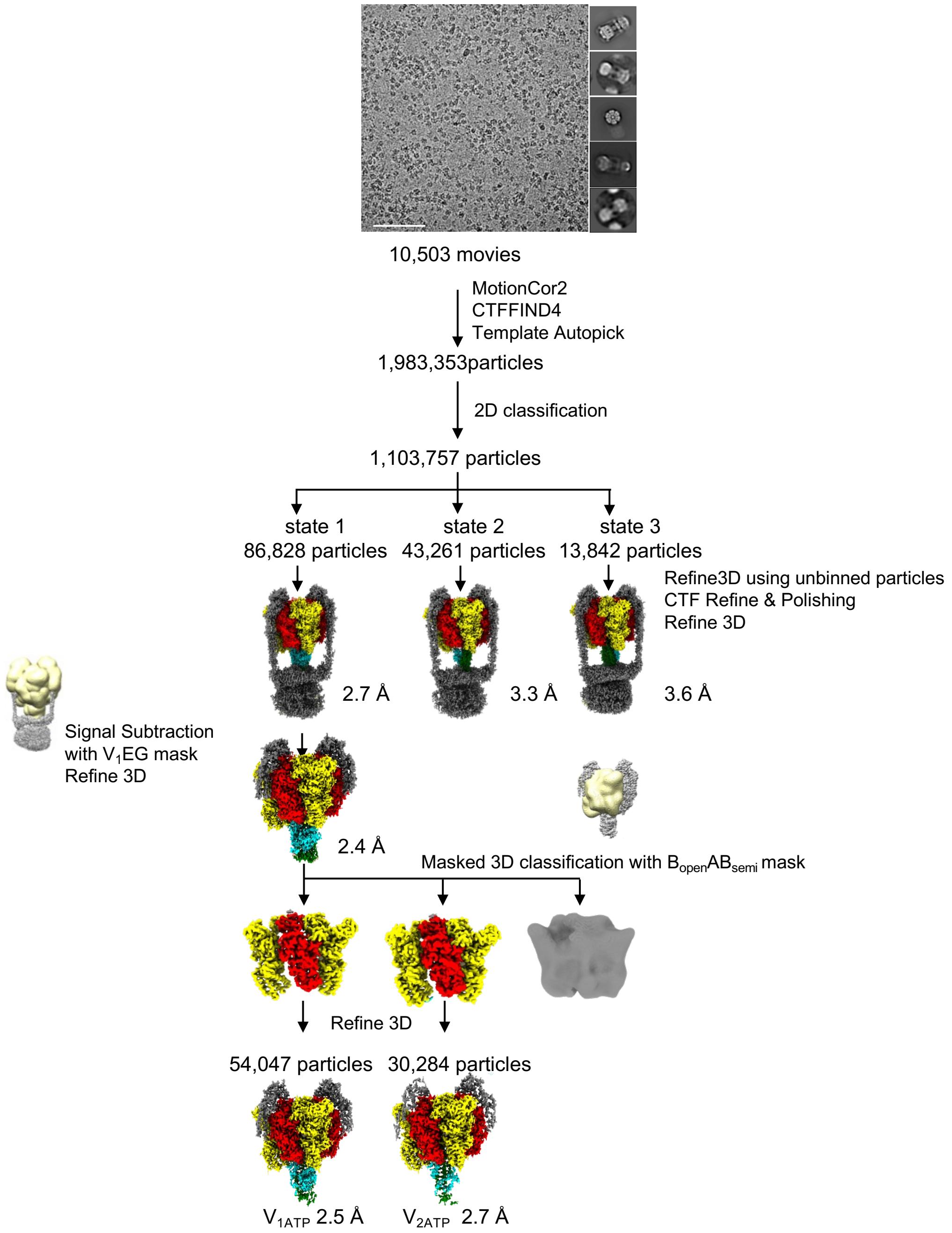

**Figure 3-figure supplement 2. Cryo-EM processing workflow of the V/A-ATPase activated by 4 mM ATP for 5 s.** A processing pipeline for 3D reconstruction of the nucleotide-free V/A-ATPase ($V_{nucfree}$) under saturating ATP concentration with 20 mM sulfate. Example micrograph, 2D class average images (top, scale bar; 100 nm), and cryo-EM data analysis workflow (bottom) are shown. Masked 3D classification allowed us to obtain two other maps with improved quality of the $V_1$EG region at 2.5 Å ($V_{1ATP}$) and 2.7 Å ($V_{2ATP}$), respectively.

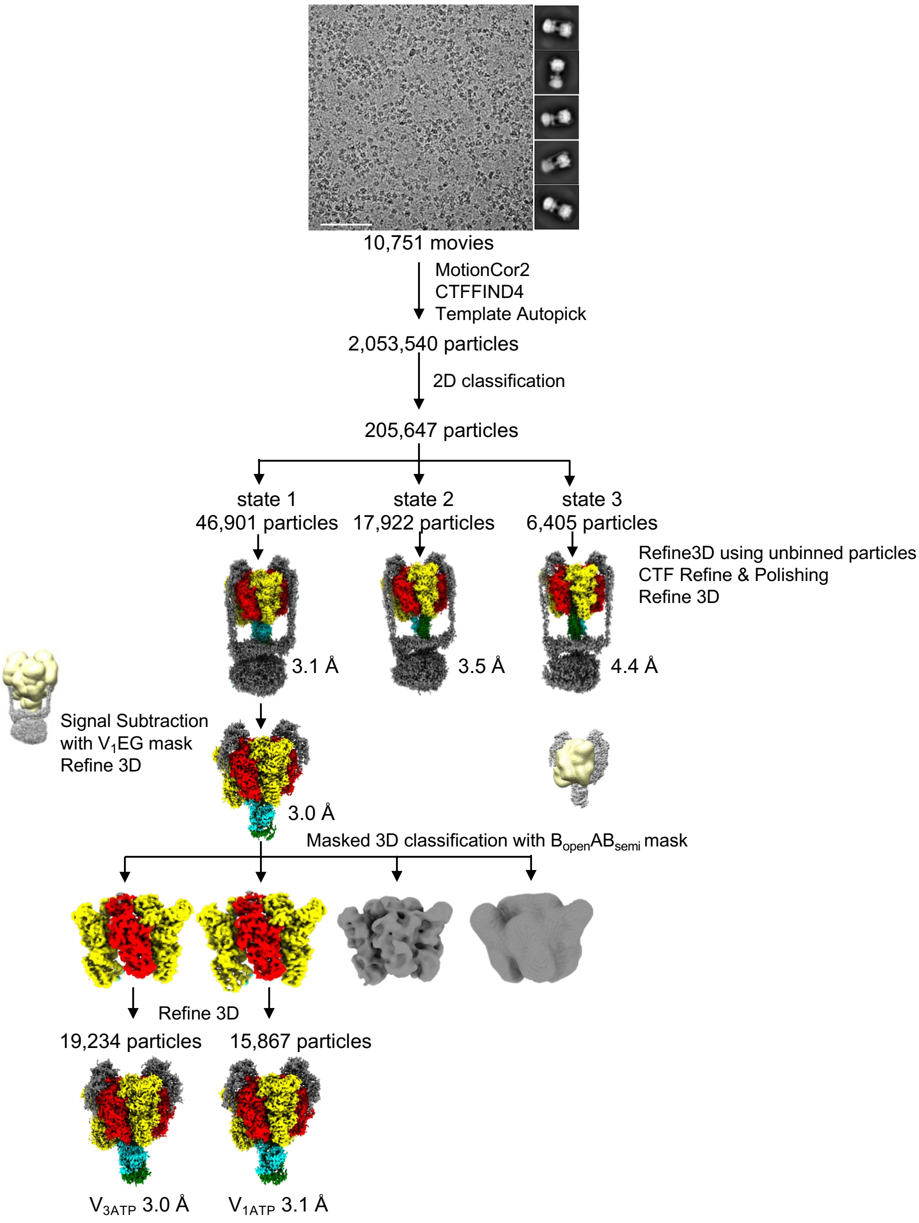

**Figure 3-figure supplement 3. Cryo-EM processing workflow of the V/A-ATPase activated by 4 mM ATP for 30 s.** A processing pipeline for 3D reconstruction of the nucleotide-free V/A-ATPase ($V_{nucfree}$) under saturating ATP concentration with 20 mM sulfate. Example micrograph, 2D class average images (top, scale bar; 100 nm), and cryo-EM data analysis workflow (bottom) are shown. Masked 3D classification allowed us to obtain two other maps with improved quality of the $V_1$EG region at 3.0 Å ($V_{3ATP}$) and 3.1 Å ($V_{1ATP}$), respectively. The $V_{1ATP}$ map was identical to the $V_{1ATP}$ map obtained from the dataset of 5 sec reaction (Figure 3-figure supplement 2); hence, a higher resolution map of $V_{1ATP}$ was used for further analysis.

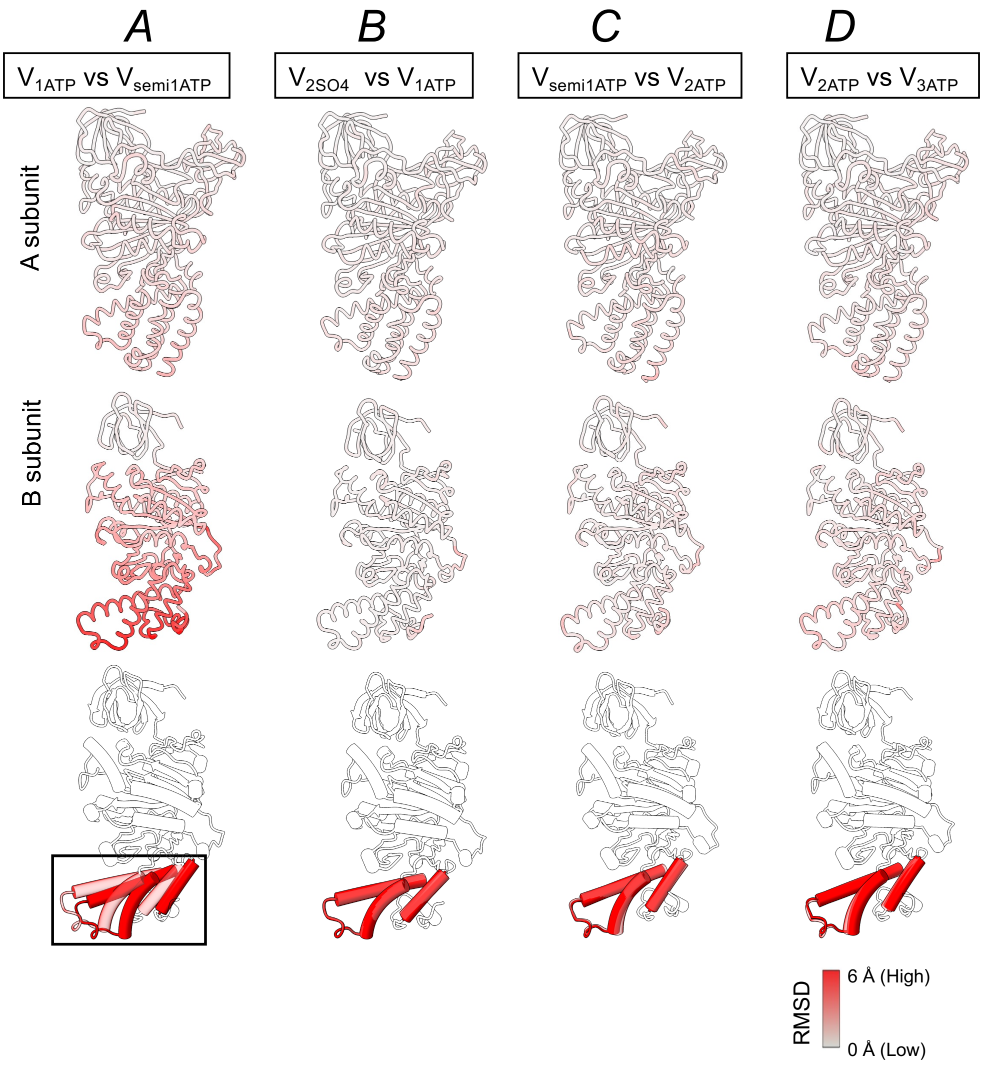

**Figure 3-figure supplement 4. Comparison between intermediates in each semi-closed A and B subunits.** The semi-closed A and B subunits of each intermediate were superimposed on the β barrel domain (A: 1–70 a.a. and B: 1–70 a.a., respectively). Models are colored by $C_\alpha$ root mean square displacement (RMSD) values calculated using $C_\alpha$ atoms; gray (small changes) to red (large changes). $C_\alpha$ RMSD values between $V_{1ATP}$ and $V_{semi1ATP}$ **(A)**, $V_{2SO4}$ and $V_{1ATP}$ **(B)**, $V_{semi1ATP}$ and $V_{2ATP}$ **(C)**, $V_{2ATP}$ and $V_{3ATP}$ **(D)** are shown in top. Comparisons of the C-terminal helix bundle of B subunit of $V_{1ATP}$ (transparent) and $V_{semi1ATP}$ (opaque) (**A**), $V_{2ATP}$ (transparent) and $V_{1ATP}$ (opaque) (**B**), $V_{semi1ATP}$ (transparent) and $V_{2ATP}$ (opaque) (**C**), $V_{2ATP}$ (transparent) and $V_{3ATP}$ (opaque) (**D**) are shown in bottom. The long α-helix of the C-terminal helix bundle of the B subunit are shown as cylinders to clarify the structural differences**.**

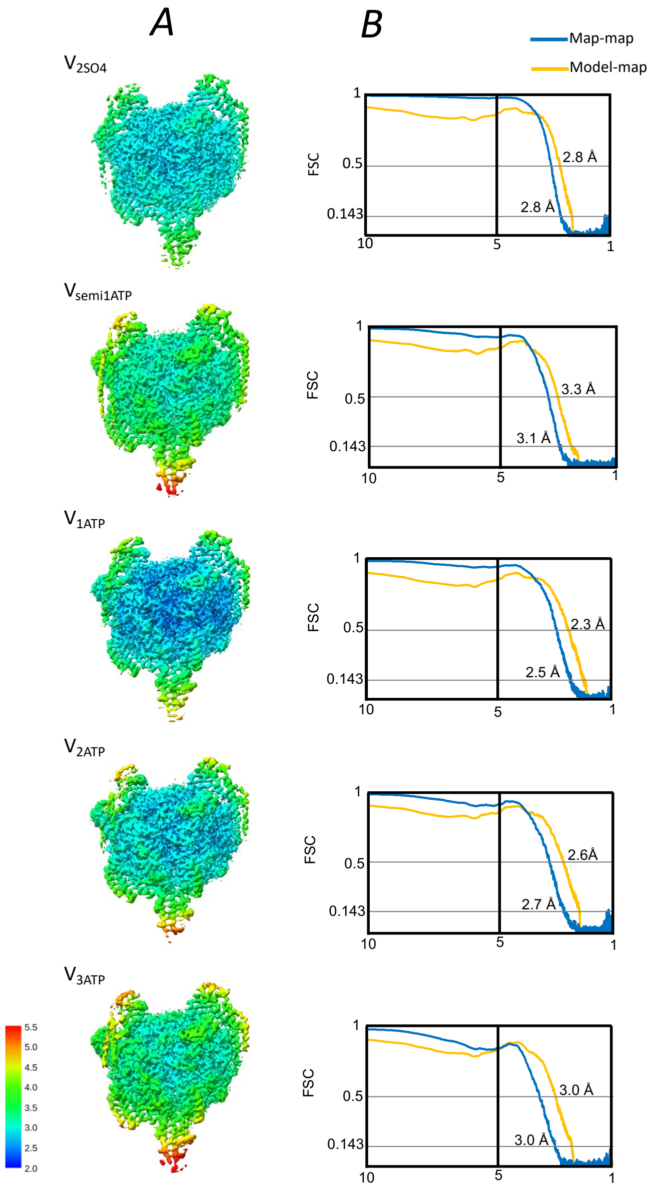

**Figure 3-figure supplement 5. Cryo-EM map validation for the $V_1EG$ domain of the intermediates.** **A**, Local resolution of the final maps of initial intermediates as indicated. Resolutions are color-coded by scale bars. **B**, Fourier shell correlation (FSC) curves of reconstructed map of $V_1EG$ domain.

| A/444-573 | $V_{2SO4}$ | $V_{semi1ATP}$ | $V_{1ATP}$ | $V_{2ATP}$ | $V_{3ATP}$ |
|---|---|---|---|---|---|
| $V_{2SO4}$ | | 0.898 | 0.380 | 1.294 | 1.917 |
| $V_{semi1ATP}$ | 0.898 | | **1.092** | 0.541 | 1.143 |
| $V_{1ATP}$ | 0.380 | **1.092** | | 1.465 | 2.083 |
| $V_{2ATP}$ | 1.294 | 0.541 | 1.465 | | 0.761 |
| $V_{3ATP}$ | 1.917 | 1.143 | 2.083 | 0.761 | |

| B/374-446 | $V_{2SO4}$ | $V_{semi1ATP}$ | $V_{1ATP}$ | $V_{2ATP}$ | $V_{3ATP}$ |
|---|---|---|---|---|---|
| $V_{2SO4}$ | | 5.11 | 0.240 | 4.813 | 5.223 |
| $V_{semi1ATP}$ | 5.111 | | **5.225** | 0.667 | 0.778 |
| $V_{1ATP}$ | 0.240 | **5.225** | | 4.923 | 5.321 |
| $V_{2ATP}$ | 4.813 | 0.667 | 4.923 | | 1.050 |
| $V_{3ATP}$ | 5.223 | 0.778 | 5.321 | 1.050 | |

**Supplementary table 1. RMSD values for C-terminal domain of semi-closed A (top) and B (bottom) subunit.** Subunits were superimposed on the β-barrel domain (1–70 a.a), and then the values for the backbone (444–573 a.a for A subunit, 374–446 a.a for B subunit) were calculated using UCSF ChimeraX. The values for $V_{semi1ATP}$ versus $V_{1ATP}$ are indicated in bold.

| | $V_{2SO4}$<br>EMD-34365<br>PDB ID 8GXY | $V_{semi1ATP}$<br>EMD-34366<br>PDB ID 8GXZ | $V_{1ATP}$<br>EMD-34362<br>PDB ID 8GXU | $V_{2ATP}$<br>EMD-34363<br>PDB ID 8GXW | $V_{3ATP}$<br>EMD-34364<br>PDB ID 8GXX |
|---|---|---|---|---|---|
| **Data collection and processing** | | | | | |
| Magnification | 60,000 | 60,000 | 105,000 | 105,000 | 105,000 |
| Voltage (kV) | 300 | 300 | 300 | 300 | 300 |
| Electron exposure (e−/Å2) | 50 | 50 | 50 | 50 | 50 |
| Defocus range (μm) | −1.2 to −2.4 | −1.2 to −2.4 | −0.8 to −1.8 | −0.8 to −1.8 | −0.8 to −1.8 |
| Pixel size (Å) | 0.81 | 0.81 | 0.83 | 0.83 | 0.83 |
| Symmetry imposed | | | C1 | | |
| Initial particle images (no.) (after 2D) | 1,452,886 | | 1,103,757 | | 205,647 |
| Final particle images (no.) | 145,880 | 39,101 | 54,047 | 30,284 | 19,234 |
| Map resolution (Å) | 2.8 | 3.1 | 2.5 | 2.7 | 3 |
| FSC threshold | | | 0.143 | | |
| **Refinement** | | | | | |
| Initial model used (PDB accession no.) | | | This study | | |
| Model resolution (Å) | 2.8 | 3.3 | 2.3 | 2.6 | 3 |
| FSC threshold | | | 0.5 | | |
| Map-sharpening *B* factor (Å2) | -76 | -67 | -41 | -39 | -67 |
| Model composition | | | | | |
| Nonhydrogen atoms | 29,472 | 29,499 | 29,503 | 29,530 | 29,553 |
| Protein residues | | | 3,788 | | |
| Ligands | $SO_4$: 2 | $SO_4$: 1, ATP: 1 | $SO_4$: 2, ATP: 1 | $SO_4$: 1, ATP: 2 | ATP: 3 |
| *B* factors (Å2) | | | | | |
| Protein | 58.51 | 91.33 | 53.2 | 70.31 | 65.86 |
| Ligand | 58.35 | 115.67 | 73.07 | 74.12 | 63.31 |
| R.m.s. deviations | | | | | |
| Bond lengths (Å) | 0.005 | 0.006 | 0.007 | 0.006 | 0.006 |
| Bond angles (°) | 0.519 | 0.535 | 0.574 | 0.546 | 0.557 |
| Validation | | | | | |
| MolProbity score | 1.3 | 1.66 | 1.44 | 1.36 | 1.49 |
| Clashscore | 5.16 | 6.61 | 4.64 | 5.14 | 5.57 |
| Rotamer outliers (%) | 0 | 0 | 0 | 0 | 0.06 |
| Ramachandran plot | | | | | |
| Favored (%) | 97.87 | 95.72 | 96.76 | 97.61 | 96.92 |
| Allowed (%) | 2.1 | 4.25 | 3.24 | 2.36 | 3.06 |
| Disallowed (%) | 0.03 | 0.03 | 0 | 0.03 | 0.03 |

**Supplementary table 2. Cryo-EM data collection, refinement, and validation statistics.**
